# Measured and modelled transitions between self-paced walking and synchronization with rhythmic auditory cues

**DOI:** 10.1101/2025.10.06.679997

**Authors:** Clémence Vandamme, Virginie Otlet, Renaud Ronsse, Frédéric Crevecoeur

## Abstract

Constraining gait rhythm with a metronome has been shown to influence gait pattern in many different ways. While rhythmic cues can improve several parameters in some clinical populations, they do alter the long-range autocorrelations naturally exhibited in series of stride durations. However, transitions between walking with and without a metronome (and vice versa) have not been measured; it is therefore unclear how people adapt to such a change in task. To address this gap, a total of 21 healthy volunteers were asked to walk overground under three conditions: one unconstrained control condition, followed by two conditions in which a metronome was activated during either the first or second half of the trial to test both transitions. The long-range autocorrelations were assessed over a sliding window on the stride series to measure their evolution. Our observations were reproduced with a computational model allowing us to relate sudden changes in movement parameters to the long-range autocorrelations, which are typically measured over longer timescales. The results showed a clear transition in both conditions involving a metronome, with long-range autocorrelations of the series of stride durations gradually reduced when the metronome was turned on and recovered when it was turned off. In these two conditions, the change in long-range autocorrelations could be reproduced in the model by an instantaneous switching of the control policy associated with the presence or not of the metronome, suggesting that long-range autocorrelations emerge from a flexible control strategy that rapidly regulates timing and amplitude parameters according to task requirements.

**Significant statement:** Through an experiment involving transitions between walking with and without a metronome, we studied how people adapt to such a change of task by measuring the evolution of long-range autocorrelations (LRA) in the stride series. The results were reproduced in a model by an instantaneous change in the control policy, which validates the hypothesis that LRA emerge from a flexible control that rapidly regulates timing and amplitude parameters according to task requirements.

## Introduction

In natural walking conditions, humans seem to have a remarkable ability to finely tune their stride-to-stride fluctuations. Despite an infinite number of pairs of stride length and duration enabling them to reach a given desired velocity, the coefficient of variation of those gait parameters is on the order of just a few percent (1). This variability has been extensively studied because it holds significant potential as a sensitive parameter for assessing mobility and fall risk in clinical settings and gaining insights into stride-to-stride control across different contexts (2). However, to fully understand how humans regulate their strides, it is necessary to analyze not only the average magnitude of the stride-to-stride variability but also the temporal structure of these fluctuations.

Indeed, gait variability in healthy adults exhibits Long-Range Autocorrelations (LRA), or statistical persistence, meaning that fluctuations at any time statistically depend on remote previous cycles with a power-law relationship, revealing the existence of a long-term dependency in the locomotor system (3). This pattern has been observed in various biological signals, such as heartbeat (4), breathing (5), neural activity (6) and finger tapping (7), but its interpretation remains debated. In gait, statistical persistence has been associated with the flexibility and adaptability of the locomotor system, while its loss has been correlated with a higher risk of falling in some pathologies such as Parkinson’s disease (8, 9). This metric is also highly sensitive to context, as evidenced by its alteration in various conditions such as cautious gait (10), forced marching in military populations and load carrying (11), walking at non-preferred velocity (12), and synchronization to external cues (13). In particular, several authors have shown that constraining gait rhythm with a metronome resulted in series of stride durations exhibiting *anti-persistence*, with little or no impact on the magnitude of this variability (13–19). In contrast to statistical persistence, which indicates that deviations from the mean are statistically more likely to be followed by deviations in the same direction, anti-persistence means that deviations in one direction are statistically more likely to be followed by deviations in the opposite direction (14).

Several models have been proposed over the years to interpret the presence of persistence in stride-to-stride variability when walking freely, and the emergence of anti-persistence in these same series when synchronizing with a metronome. While many authors consider that LRA in stride interval naturally arise from the complex interaction between different structures acting at different time scales (20), others suggested that it might be due instead to the inherent biomechanics of walking and may not require complex central nervous system control mechanisms (21). It has also been argued that LRA might simply be a predictable consequence of partial error correction (22). In that sense, Dingwell and colleagues (23) demonstrated that when walking on a treadmill, humans do not immediately correct deviations in stride duration and length, as these two parameters are irrelevant to the primary objective of the task, namely maintaining a stable velocity. In contrast, they showed that humans would slightly overcorrect small deviations in stride speed, resulting in slight anti-persistence in series of stride velocities.

Finally, the use of rhythmic cues has also been extensively studied in populations suffering from gait impairments, such as Parkinson’s disease, especially for rehabilitation purposes. It has been hypothesized that a metronome effectively improves stride length, cadence and speed, and reduces stride-to-stride variability (24). However, as stated earlier, invariant cueing disrupts the natural statistical persistence of stride-to-stride fluctuations. To address this limitation, there is a growing interest in fractal metronomes, which would reinstate a pattern of variability similar to that of healthy individuals (25). In this context, few authors investigated whether the benefits of rhythmic cues, either visual or auditory, isochronous or fractal, could be retained, with promising results in the elderly and in patients with Parkinson’s disease (26–28). Nevertheless, in these studies, the temporal characterization of the retention remained poor, consisting of a single measurement of the Hurst exponent over a post-synchronization phase. There is therefore a need for new methods to analyze the transition between phases with and without rhythmic cueing and to understand the effect of the metronome better.

To address this gap, we investigated the impact of the sudden activation or deactivation of an isochronous metronome by measuring the evolution of the (anti-)persistence of new experimental stride series over a sliding window. Under the model linking the presence of persistence to goal-directed correction (23), we reproduced in theory the effect of the metronome and tested different changes in control policies at the transitions. Our observations were compatible with an instantaneous adjustment of the control policy in the model, resulting in a modulation of LRA associated with the synchronization of the cadence with the metronome. Together, our results highlight a functional link between statistical persistence and a flexible modulation of gait control, and suggest that healthy adults rapidly and efficiently adapted their gait to task requirements.

## Materials and methods

### Experimental procedure

A total of 21 young adults (9 females, age: 25, 4 ± 1.5 yrs; height: 175.3 ± 11.8 cm; mass: 75.2 ± 14.3 kg) were enrolled in this study and performed the three conditions described below. The target sample size was based on previous studies reporting a significant effect of an isochronous cueing on statistical persistence (13–15, 17, 18). Participants were naive to the purpose of the experiment and had no known neurologic, muscular, or orthopedic disorder that could alter their gait. The ethics committee of the host institution (UCLouvain) approved the experimental procedures, and participants provided written informed consent prior to the experiment, in accordance with usual procedures.

Each participant walked along a 34-meter-long rectangular track with rounded corners for 15 minutes in three conditions: a first control condition without a metronome to measure their spontaneous cadence, followed by two conditions involving a transition between a phase without a metronome, and a phase with a metronome imposing the cadence of their gait (Fig. 1A), or vice-versa. In these two conditions, the metronome was activated either during the first half of the trial (ON-OFF condition) or during the second half of the trial (OFF-ON condition). Participants were instructed to synchronize one foot to the metronome beat. The order of the conditions involving a metronome was counterbalanced across the participants. The metronome was set to the frequency corresponding to the participants’ spontaneous cadence, recorded as the average cadence during the control condition. The cueing was isochronous, meaning that the inter-beat intervals did not display any variability. During a training lap with the metronome, the frequency was tested and adapted, with a maximum adjustment of ± 5% when requested by the participant. Participants were asked to maintain a constant speed within each trial and were explicitly instructed not to interrupt their walk when the metronome was switched on or off.

**Figure 1:**
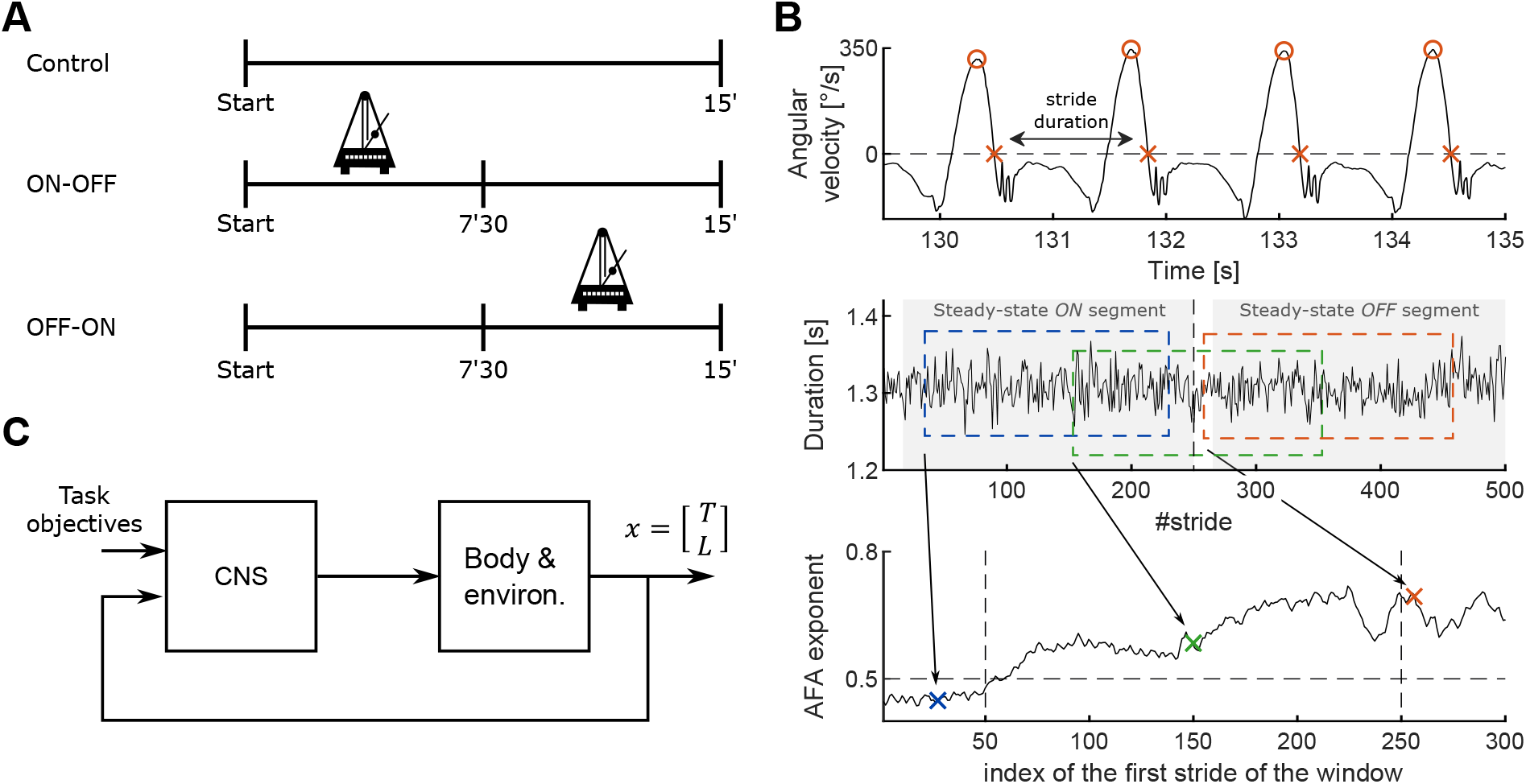
Experimental paradigm, data processing and model of stride-to-stride regulation. (A) Overview of the 3 conditions completed by the participants, with durations and periods when the metronome was on. (B) Top to bottom: illustration of stride duration derivation from the sagittal angular velocity recorded by an IMU placed on the participant’s ankle, example of series of stride durations in control condition and evaluation of the evolution of the AFA exponent over a sliding window. (C) Schematic representation of the model of gait control.

### Data acquisition and processing

Data were collected using NGIMU (x-io Technologies, Bristol, UK), and MATLAB R2021a (The MathWorks, Natick, MA, USA) was used for data processing, analysis, and statistics (“Statistics and Machine Learning Toolbox”). Participants wore four Inertial Measurement Units (IMUs), placed on both ankles (just above the lateral malleolus) and on top of both feet. These IMUs were used to obtain the sagittal angular velocities at a sample rate of 400Hz, and derive series of stride durations. Stride durations were defined as the time between two consecutive heel strikes of the same foot, calculated from the detection of the zero crossings following each peak in the sagittal angular velocity (Fig. 1B, top panel) (29). By default, the IMU used to extract the stride durations was the one attached to the left ankle. However, due to connection losses between the IMUs and the computer, some trials displayed gaps in the recorded data. In those cases, we used the IMU attached to the right ankle or those placed on the feet. When IMUs from different sides could be used to reconstruct stride series, the properties of the resulting stride series were similar (mean, standard deviation, and autocorrelation functions). Therefore, the IMU used for analysis had no impact on the results. For some trials, connection losses of the two IMUs did not allow us to reconstruct the stride series with sufficient accuracy (more than 10% missing strides). Such a number of missing strides could be critical for characterizing the temporal structure of the variability and could prevent a proper computation of the AFA exponent. Four trials of different participants were therefore discarded from the analyses (three trials in the control condition and one trial in the ON-OFF condition). The remaining trials had a maximum of 2% missing strides, which should have no impact on the evaluation of the statistical persistence, according to several studies (30, 31). All trials in the control conditions included enough strides to estimate the average pace of the participants and determine the metronome frequency.

In addition, four more trials were discarded because these participants clearly demonstrated an inability to perform the task, i.e., to synchronize one foot with the metronome beat. Besides the difficulties reported by these participants themselves during the experiment, these series exhibited large oscillations around the instructed pace. These oscillations were of a similar or even greater magnitude than in the non-paced condition, showing that they were unable to adapt their cadence to the target value and follow the instruction. To determine which trials needed to be excluded, we plotted a moving average with a window of 50 data points in the ON part of the series (ON-OFF and OFF-ON trials) and calculated the Root Mean Squared Error (RMSE) from the metronome instruction. Outliers were defined as trials with RMSE 1,5 times the interquartile range above the third quartile (*RMSE* > *Q*_3_ + 1.5 *IQR*). The four series that were removed based on this criterion were the two trials with the metronome of one participant, and the OFF-ON condition of two other participants. These series had an RMSE greater than 0.0085*s*, whereas the series retained for analysis had an RMSE below 0.0041*s*. In sum, 55 series out of the 63 recorded ones were retained for analysis.

To confirm that participants adopted a stable speed, we derived an estimate of stride lengths in the control condition, based on the method described in (32). We computed the stride velocities by dividing these lengths by the corresponding stride durations and computed the range of variation of each series of stride velocities, smoothed with a moving average of 10 strides. We reported a maximum range of variation of 15% across all the participants in the control condition.

### Data analysis

In the ON-OFF and OFF-ON conditions, we decomposed the stride series into two parts to compare the series with and without the metronome in a time window that did not involve the transition. The first 15 strides of the pre-switch and post-switch series were discarded in order to restrict our analysis to steady-state behavior and avoid biases that may arise from non-stationarities. These steady-state segments are highlighted by the shaded areas in Fig. 1B, middle panel. These resulting series ranged from 340 to 420 strides according to each participant’s pace. We estimated the standard deviation and the level of LRA of these series to assess the behavioral changes induced by the metronome.

The level of LRA was computed using the Adaptive Fractal Analysis (AFA), a method described in (33) and recommended for estimating fractal exponents of stride time series due to its robustness to non-stationarities (34, 35). Briefly, the AFA consists of identifying smooth trend signals from a time series. These signals are derived from a linear combination of local polynomial fits applied to overlapping windows of a fixed size. The process is repeated for window sizes ranging from 11 strides to half the length of the series, evenly spaced on a logarithmic scale. The residuals of each fit are reported against the corresponding timescale, and the analysis of the scaling of the residuals as a function of the timescales gives the *AFA exponent* of the time series. This exponent estimates the same quantity as the exponent obtained with the Detrended Fluctuation Analysis (DFA), also known as *α* exponent or Hurst exponent (*H*), which captures randomness or (anti-) persistence in the time series. When 0 < *H* < 0.5, the series is anti-persistent. When *H* = 0.5, the series corresponds to random fluctuations around its mean (white noise), and when *H* > 0.5, the series displays statistical persistence associated with LRA. The Hurst exponent is by definition restricted within the interval]0,1[while the AFA can yield values higher than 1. This corresponds to non-stationary series, a case that was not encountered in the present experiment.

To characterise the behavioral response following the change in task instruction set by turning ON or OFF the metronome, we then analyzed the evolution of the AFA exponent by applying the AFA over a sliding window of 200 strides involving the transition (Fig. 1B). All strides were considered for this analysis, strides excluded from the steady-state segment analysis were re-integrated to interpret the transition correctly. We aimed to use the smallest possible windows to characterize the transitions over the shortest time scale. However, windows of size below 200 strides resulted in noisy profiles, altering the interpretability of the results. While numerous studies recommended using DFA on series of at least 500 strides (36–38), evenly-spaced regressions, as employed in this study, allow reducing the required number of strides to approximately 200 by reducing the sensitivity of the slope estimation to data variability (39). Other authors suggested that the AFA can be used to estimate the Hurst exponent for time series of approximately 200 strides (40), further supporting the choice of window size in the present study.

In the conditions involving a metronome, some series displayed a discontinuity at the transition, i.e., a sudden increase or decrease in the mean stride durations, likely resulting from short-term adjustments to the introduction or removal of the metronome. Such an abrupt change could artificially increase the AFA exponent as long as the discontinuity falls within the sliding window, as it introduces an artificial transient effect that may lead to erroneous conclusions about changes in LRA. To address this potential confounding factor, we applied a correction to remove this discontinuity for all trials in ON-OFF and OFF-ON conditions. Then we compared the evolution of the AFA exponents with and without the correction. This correction consisted of subtracting an offset from the second part of the series, which corresponded to the difference between the average value of the first 15 strides after the switch and the average value of the last 15 strides before the switch. In addition, the same offset measured in each trial was used to build surrogate data to perform a control analysis, as explained below. Surrogate data are often used in gait analysis to assess whether observed variability patterns truly reflect non-linear dynamics (41). We adapted this technique to distinguish the effect of a potential change in average pace from a change in experimental constraints. For each trial in the ON-OFF condition, we extracted the ON part of the series and duplicated it. We then added the offset to the duplicated series and concatenated it with the other, thus obtaining artificial “ON-ON” series reproducing the same discontinuity encountered experimentally, but without any change in task. For each trial in the OFF-ON condition, we followed the same procedure to obtain “OFF-OFF” series. In both cases, the surrogate series allowed us to assess the effect of a short-term change in the mean of stride series, which could be dissociated from the change in gait pattern corresponding to the metronome.

### Model

Stride series with a similar evolution of LRA as those obtained experimentally were then reproduced within the framework of stochastic optimal control (42). This framework offers an interpretation of the transitions as task-dependent changes in gait control, mediated by the introduction or removal of the metronome. The model developed here was based on the study by Dingwell and colleagues (23)(43), and adapted to reproduce a broader range of behavior, such as the anti-persistence emerging from metronome-guided walking, as well as transitions between these two tasks. A schematic representation of the model is provided in Fig. 1C. Briefly, each stride was generated iteratively from the previous one according to the following discrete-time system:

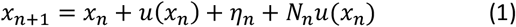

Each stride was characterized by a state vector, *x*_*n*_, composed of two state variables defined as the stride duration and length (*x*_*n*_ = [*T*_*n*_, *L*_*n*_]^*T*^). The noise was composed of an additive noise vector *η*_*n*_ and a command-dependent noise *N*_*n*_*u*(*x*_*n*_). The vector *η* and the diagonal of the matrix *N* were zero-mean, Gaussian random variables with covariance matrix equal to 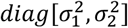, off-diagonal terms being equal to 0. The command *u*(*x*_*n*_) can be seen as the stride-by-stride regulation of length and cadence against noise disturbances according to the behavioral goal.

The initial formulation of the model was designed to reproduce treadmill walking, without a metronome (23). Therefore, the overall objective was to maintain a constant velocity. As an infinite amount of [*T*_*n*_, *L*_*n*_]^*T*^ allows maintaining a constant velocity, there is one degree of freedom in the set of state variables for preserving a constant speed, making the task *redundant*. However, along this redundant axis, one pair, called the Preferred Operating Point (POP) and denoted as [*T*_*POP*_, *L*_*POP*_], may be naturally preferred, likely due to biomechanical constraints on the achievable range of cadence and length modulation. A secondary objective was then to limit deviations from this preferred pair of stride length and duration. These two objectives were balanced with the energetic cost that this regulation required. The cost function (Eq. 2), used to compute the motor command, captures these objectives, each one weighted by a factor (*α, β, γ*) reflecting its relative importance:

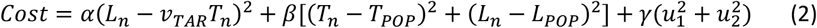

The first term penalizes deviations from the targeted velocity *v*_*TAR*_; the second term, weighted by *β*, penalizes deviations away from the POP, and the last term weights the cost of the command. Finding the optimal command is a classical Linear-Quadratic-Gaussian (LQG) control problem, the derivation of which is described in detail in (44). The command is optimal in the sense that it minimizes the expected value of the quadratic cost criterion defined by Eq. 2, and the solution is a linear function of the state (optimal feedback gains). As we aimed to model steady-state walking, both the system and the cost were time-invariant. Therefore, we used the infinite horizon formulation. The optimal state feedback gains were time-invariant and were obtained by solving the Algebraic Riccati Equation (42).

The parameters of this model were adjusted to match the level of variability of the series of stride durations and its temporal structure. In the control condition, i.e., to model overground walking without a metronome, the weights of the cost function were set to *α* = *β* = 2.5 and *γ* = 10. The penalty applied to deviations from *T*_*POP*_ and *L*_*POP*_ was therefore relatively small compared to the weight applied to the cost of the motor command, allowing deviations away from the POP to persist, and the series of stride durations and lengths to exhibit fractal-like fluctuations, i.e., LRA. Here, the penalty applied to deviations from the velocity *v*_*TAR*_ was also set to 2.5 to account for the fact that the velocity was not strictly regulated since our participants were not walking on a treadmill. To simulate treadmill walking, this parameter would be considerably higher. The target velocity was 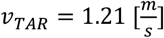, with *T*_*POP*_ = 1.189 [*s*] and *L*_*POP*_ = *v*_*TAR*_*T*_*POP*_ [*m*], which was set to the mean velocity and stride duration adopted by the participants.

The model was then adapted to reproduce the properties of stride series when participants were asked to synchronize with a metronome. Our initial approach was based on modulating the parameter *β* to simulate stronger regulation around the POP. However, we observed that this approach did not fully explain the behavior. In fact, to synchronize with a metronome, participants needed to regulate the asynchronies between the heel strike of one foot and the metronome beat, or in other words, adapt the stride duration they targeted at each stride. If one stride felt behind the metronome, the following stride needed to be shorter to catch up with the metronome. In the model, a target duration 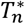 was iteratively computed as the duration between two beats of the metronome (fixed, noted *T*_*met*_) added to the previous stride error. The initial system dynamics (Eq. 1) were augmented to include this supplementary state variable:

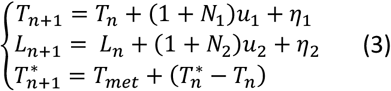

The cost function was also adapted to include a term penalizing errors on the target stride duration 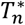 at each time step:

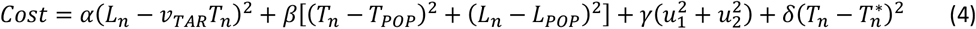

The parameter weighting the synchronization to the metronome was set to *δ* = 0.35 in order to fit the average AFA exponent observed experimentally when the metronome was turned on. The duration of the metronome *T*_*met*_ was set to *T*_*POP*_ ≔ 1.189*s*, as the metronome was set at the frequency corresponding to the participants’ spontaneous average cadence in the experimental setting.

Different transitions from *δ* = 0 to *δ* = 0.35 (and vice-versa) were tested. The transitions were initiated at different rates: infinite slope, corresponding to instantaneous transitions; or slopes set to 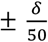, or 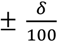). Negative (resp. positive) slopes correspond to a decrease (resp. increase) of the regulation to the metronome beat and therefore refer to the ON-OFF (resp. OFF-ON) condition. The controller was then recomputed following changes in *δ* to adapt the feedback gains. Each set of parameters was used to generate 100 series in the ON-OFF and OFF-ON conditions. We used the onset of change in LRA to identify the change in *δ* that matched the experimental observations.

We also varied the initial asynchrony by manipulating *T*^∗^ at the transition of the OFF-ON condition. The initial value of *T*^∗^could be set to *T*_*met*_, capturing a perfect synchronization of the first beat of the metronome with the corresponding stride. To match the experimental settings more realistically, the initial asynchrony was also randomly drawn from a uniform distribution 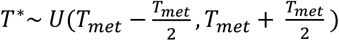. These boundaries were chosen to match the worst initial asynchrony that participants could experience, which corresponds to the case where the metronome beat occurred exactly between two strides. In the ON-OFF condition, the metronome was activated before gait initiation. We therefore assumed an initial asynchrony of zero (*T*^∗^ = *T*_*met*_).

The parameters (transition rate and initial condition) were selected to reproduce the evolution of the AFA exponents observed experimentally. All statistical analyses were then applied to a sample of 20 series generated with the selected parameters, similarly to the experimental data. All model parameters and the criteria for choosing their values are summarized in Table 1.

**Table 1.** Model parameters.

| Parameters | Criteria |
| --- | --- |
| $\sigma_1^2 = 0.017 [s^2], \sigma_2^2 = 0.021[m^2]$ | Values from (23) |
| $\alpha = \beta = 2.5$<br>$\gamma = 10$ | Fit to variability and AFA exponent of series of stride durations in control condition. |
| $\delta = 0$ if metronome OFF<br>$\delta = 0.35$ if metronome ON | Fit to global AFA exponent of series of stride durations when metronome was activated. |
| $v_{TAR} = 1.21 [m/s], T_{POP} = 1.189 [s]$<br>$L_{POP} = v_{TAR} \cdot T_{POP}$ | Average values obtained from the control condition |
| $T_{met} = T_{POP}$<br>$T_{init}^* \sim U(T_{met} - \frac{T_{met}}{2}, T_{met} + \frac{T_{met}}{2})$ | Experimental protocol |
| Transition rates from $\delta = 0$ to $\delta = 0.35$ | Fit to the onset of change in AFA exponent whose evolution was outlined using a sliding window. |

Finally, although this variable was not measured in the protocol, the model allowed us to extract the series of synchronization errors, or asynchronies, defined as the time interval between heel strikes and metronome beats, when activated. This series was noted *A*_*n*_ and computed as follows: 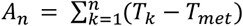, with *T*_*k*_ the duration of stride *k* and *T*_*met*_ a constant denoting the duration between two beats of the metronome. We computed the average AFA exponent and standard deviation of these series, to compare them with values previously reported in the literature (13, 18).

### Statistical analyses

We performed one-tailed paired t-tests to test the specific hypothesis, derived from prior work (13–15, 17, 18), that both the AFA exponent and the standard deviation of series of stride durations would be lower during the ON parts than during the OFF parts of the series. Effect sizes were characterized with Cohen’s d, defined as the difference between the means divided by the pooled standard deviation.

## Results

### Behavioural experiments

We first analyzed separately the ON and OFF steady-state segments of the strides series and reported the AFA exponents and standard deviations of all participants in both conditions including the metronome in Fig. 2. As expected, we observed lower AFA exponents when participants were guided by the metronome (Fig. 2A). The strides series exhibited anti-persistence with an average AFA exponent significantly lower than 0.5 (AFA exponent = 0.42 ± 0.11, p = 0.003, in the ON-OFF condition). In contrast, the average AFA exponent of the OFF parts of the series was equal to 0.79 ± 0.10, which is commonly reported in the literature regarding free walking in healthy adults (3, 45). We also observed an effect of the metronome on the standard deviation of series of stride durations, associated with a decrease when the metronome was turned on (Fig. 2B). We performed one-tailed paired t-test to evaluate whether the observed differences were statistically significant. Standard deviation of stride series significantly increased when the metronome was muted compared to series of strides set to the rhythm of the metronome in the ON-OFF condition (Δ*std* = 8.3 ∗ 10^−4^ ± 0.0019 [*s*], *t*(18) = 1.95, *p* = 0.033). A statistical difference was also observed in the OFF-ON condition, where the standard deviation decreased after turning on the metronome (Δ*std* = −0.0015 ± 0.0035 [*s*], *t*(17) = −1.79, *p* = 0.045). Although the p-values were just below 0.05, this indicated a clear trend. The effect size, as measured by Cohen’s d, was *d* = 0.22 in the ON-OFF condition and *d* = −0.38 in the OFF-ON condition, indicating a small effect. Regarding the autocorrelation functions, the AFA exponent of series of stride durations significantly increased when the metronome was not activated in both conditions (ON-OFF: Δ*AFA* = 0.34 ± 0.15, *t*(18) = 9.97, *p* < 10^−4^; OFF-ON: Δ*AFA* = −0.33 ± 0.17, *t*(17) = −7.93, *p* < 10^−4^). In contrast to the standard deviation, the effect was very strong, with *d* = 3 in the ON-OFF condition and *d* = −2.3 in the OFF-ON condition (Fig. 2).

**Figure 2:**
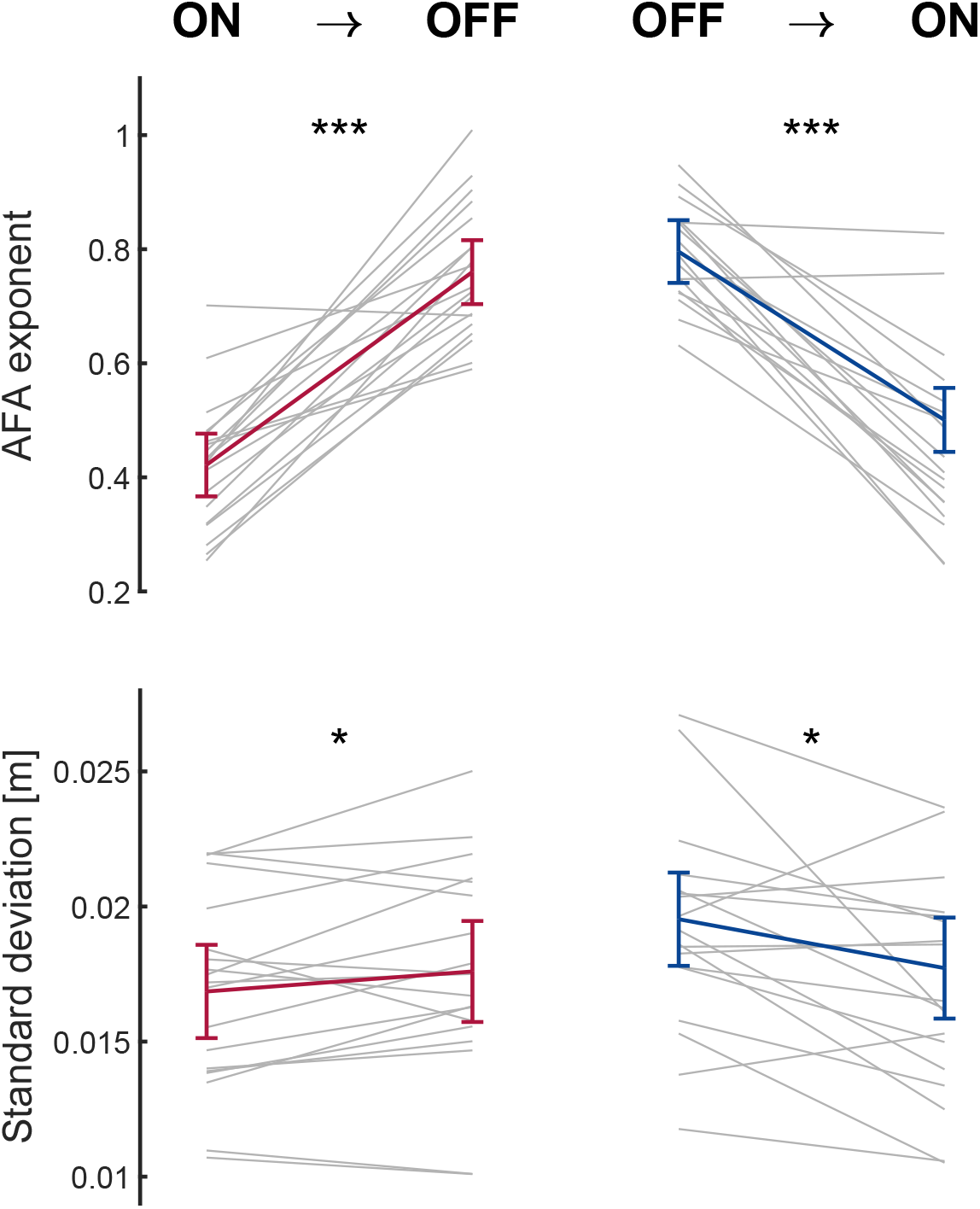
Analysis of differences between ON and OFF stride series in behavioral data. AFA exponent (A) and standard deviation (B) of series of stride durations before and after the switch of the metronome (shaded areas in Fig. 1B), in ON-OFF condition (left) and OFF-ON condition (right). Group average and 95% confidence interval are shown in color and individual participants data are shown in gray lines. Stars indicate significant differences determined using paired, one-tail t-test: * *p* < 0.05, *** *p* < 0.001.

We then investigated how participants switched from one task to another. Fig. 3A shows an example of a series of stride durations for each condition, all from the same participant. In the middle (resp. right) panel, the vertical dotted line represents the stride at which the metronome was switched off (resp. switched on). The evolution of the AFA exponent of each series was estimated by applying the AFA over a sliding window of 200 strides (see Methods). On average, participants took 765 ± 57 strides per condition, depending on the pace they chose to adopt. Therefore, to compare and average the evolution of LRA of all participants, the series were all aligned to the stride at which the metronome was turned off or on, and truncated to the same length. The evolutions of the AFA exponent of the stride series of each participant are represented in Fig. 3B, with one highlighted series corresponding to the representative participant presented in Fig. 3A. In the conditions involving a transition, the two vertical lines indicate the interval of AFA exponents measured on windows containing strides on either side of the transition. In the control condition, the AFA exponent remained relatively stable, despite clear fluctuations. Regarding the exemplar participant, we observed a clear increase in the AFA exponent in the ON-OFF condition, from a level corresponding to white noise (AFA exponent = 0.5) to a level similar to that of the control condition (AFA exponent > 0.5). It can be observed already that the AFA exponent started increasing directly after the first dotted line (i.e., as soon as the time window included the first stride from the OFF condition), suggesting that the participant recovered LRA quickly after the metronome was turned off. Conversely, the AFA exponent was gradually reduced after switching on the metronome in the OFF-ON condition, and reached a level lower than 0.5, corresponding to anti-persistence. The presence of high variability across participants and intrinsic fluctuations in the AFA exponent of each participant can be observed from the gray curves, which represent each participant. Although there was variability across participants, the average trend among all participants was reliable and similar to that of the representative participant (Fig. 3C).

**Figure 3:**
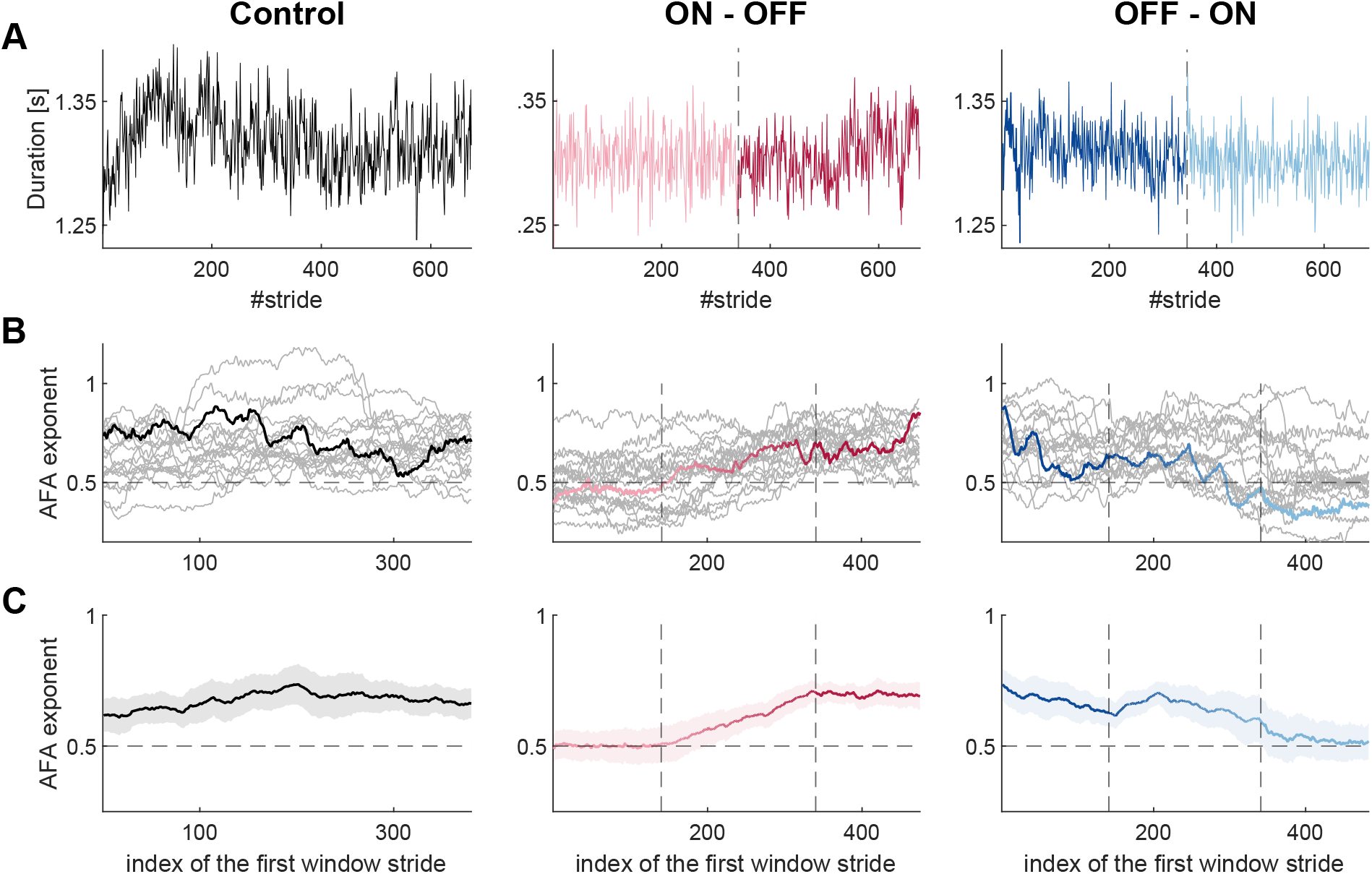
Transition analysis in behavioral data. (A) Series of stride duration for one exemplar participant in three conditions: control (left), ON-OFF (middle) and OFF-ON (right). Data in light shades (resp. dark shades) correspond to strides performed with (resp. without) the metronome, the dotted vertical line corresponds to the metronome switch. (B) Evolution of the AFA exponent of the stride series of each participant estimated by applying the AFA over a sliding window of 200 strides. Traces in color correspond to the series of the exemplar participant presented above. The two vertical lines indicate the interval of AFA exponents measured on windows containing strides on either side of the transition. (C) Same as (B) averaged over all participants with 95% confidence interval shaded area).

On average, the AFA exponent of series of stride durations in the control condition displayed a consistent trend and stabilized around a value of 0.67 (Fig. 2C, left panel). The standard deviation of individual traces ranged from 0.05 to 0.18. In the ON-OFF condition, the mean level of LRA of the series of stride durations evolved from a stable AFA exponent of 0.5 (with individual SD from 0.017 to 0.096) to a level similar to the control condition (i.e., *AFA exponent* ≅ 0.69, range of individual *SD* = [0.024, 0.10]). The differences in AFA exponents between the ON and OFF parts were smaller than those shown in Fig. 2A, likely due to a bias in the AFA when applied to shorter stride series corresponding to the moving window. Regarding the transition, the AFA exponent increased linearly as the AFA was applied to windows advancing after the switching off, such that the series contained increasingly more strides from the OFF phase. To determine the onset of this increase, we performed a segmented linear regression (46). The first segment fitted data from the first AFA exponent of the average trace (*x*_1_) to the sample *x*_*i*_, restricted to a slope of 0. The second segment fitted data from *x*_*i*_ to *x*_200_, corresponding to the stride window with an AFA exponent significantly larger than 0.5, according to the following criteria: *mean*(*AFA exponent*) − 2 ∗ *SEM* > 0.5. We then selected *x*_*i*_ such that the sum of the residuals of both fits was minimized. This analysis confirmed that the AFA exponent started to increase as soon as the window contained only three strides of the OFF phase, i.e., three points after the first vertical line in Fig. 3C, middle panel.

Similar to the ON-OFF condition, we observed a transition between two steady states in the OFF-ON condition, from a level of LRA corresponding to the control condition (*AFA exponent* = 0.67, range of individual *SD* = [0.017, 0.12]), to a level of LRA corresponding to white noise (*AFA exponent* = 0.53, range of individual *SD* = [0.021, 0.16]). In contrast to the ON-OFF condition, the AFA exponent did not decrease instantly after switching off the metronome; instead, it even slightly increased just after the transition before decreasing.

To ensure a proper interpretation of these transitions, we considered the possibility that the discontinuity in the series, occurring when the metronome was switched on or off, artificially increased the AFA exponent. An upper bound for this increase was estimated using surrogate series that featured an abrupt change in mean value of the same magnitude as the experimental data (see Methods). The AFA exponent of the surrogate ON-ON series reached an average value of 0.547 ± 0.014 when the AFA was applied to stride windows containing the discontinuity (Fig. 4A, yellow trace, between the vertical bars), while the value was on average 0.501 ± 0.004 otherwise. Regarding the OFF-OFF series, we also observed an increase in the AFA exponent for windows including the transition (mean AFA exponent = 0.789 ± 0.044) compared to stride windows without any transition (0.673 ± 0.026), of even greater magnitude (Fig. 4B, yellow). This was explained by the more pronounced discontinuities in the OFF-ON condition, which was due to the nature of the task. The sudden need to synchronize with the metronome’s rhythm required an abrupt change if the pace adopted before the transition differed from the metronome’s pace. In contrast, if the metronome’s cadence did not precisely align with the participant’s natural rhythm, they could recover that natural cadence more gradually once the metronome was turned off. These results motivated the application of an adjustable offset to the second part of each series of stride duration (after metronome transition) to mitigate these discontinuities. The average evolutions of the AFA exponent of these series (i.e., after correction) were already presented in Fig. 3C and are represented in Fig. 4A-B (black traces) together with the evolution of the AFA exponent of the initial series (gray traces). Together, these results further refined the interpretation of a rapid increase after the transition, particularly in the case of the ON-OFF transition, where the onset of the AFA exponent increase is crucial. While the presence of discontinuities indeed impacted the AFA exponent, ON-ON surrogates revealed that this effect remained limited in this condition. In addition, we still observed a rapid increase in the AFA exponent after correction, consolidating the assumption that this rapid increase reflected a genuine change in control strategy.

**Figure 4:**
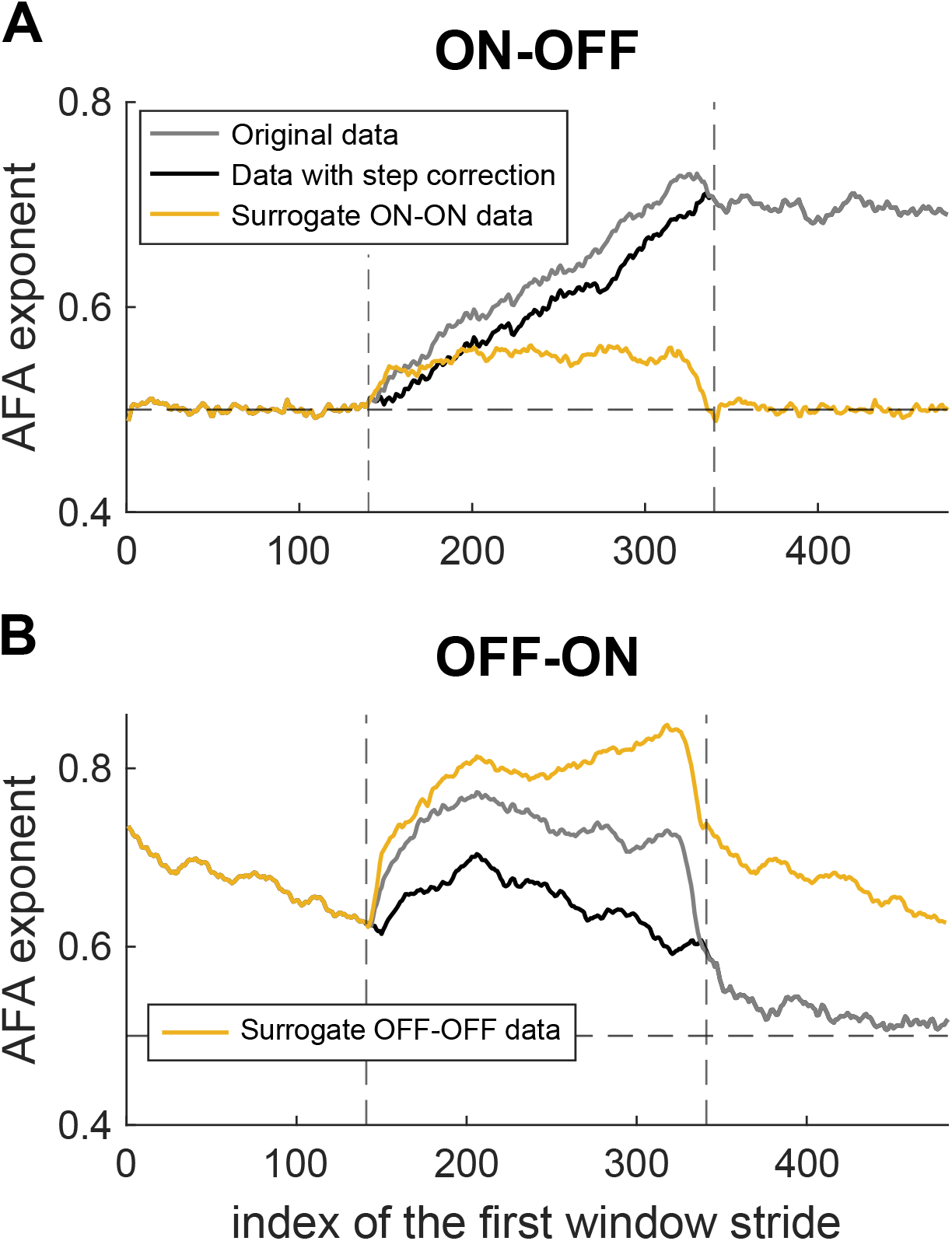
Analysis of the impact of discontinuities in stride series on the AFA exponent. Mean evolution of the AFA exponent of stride series before (gray) and after (black) applying an offset to the second part of the series to reduce the discontinuities (i.e. difference in mean value at the metronome shift). In yellow, mean evolution of the AFA exponent of surrogate data built to reproduce the discontinuities without any change of task (ON-ON and OFF-OFF series).

Collectively, the behavioural results highlighted a rapid transition of the AFA exponent between the values corresponding to ON and OFF conditions measured separately. These transitions were more pronounced when measured based on the AFA compared to the series standard deviation (and after controlling for average offsets), suggesting that LRA capture subtle changes in gait regulation that may be more difficult to identify based on standard metrics. However, since the AFA exponents were evaluated on series of 200 strides, it was difficult to precisely establish the number of strides required to adapt to the removal/addition of the metronome. The seemingly gradual change in the AFA exponent was, in fact, a consequence of the method used, consisting of applying the AFA to windows gradually containing more and more strides that occurred after the switch of the metronome. To assess the transitions in more detail, it was necessary to replicate them in a computational model that allowed us to simulate and test different hypotheses. Indeed, the almost instantaneous increase observed during the transition from ON to OFF condition, as soon as the time window included data from the OFF conditions, suggested that participants’ sensorimotor control of gait could change rapidly. Our modelling work confirmed this hypothesis.

### Model

Using the model, we tested different transitions in the control policy and assessed their consequences on the measured AFA exponents, applying the same approach to both simulated and recorded series. We simulated strides series using different candidate parameters to investigate how we could reproduce participants’ behavior. Our basic premise was that gait features were flexibly regulated, allowing for fluctuations in stride duration and length in the absence of a metronome. In contrast, a control strategy with stricter regulation of stride duration was applied with a metronome, to comply with the task instructions. A target duration (*T*^∗^) was iteratively computed for each stride to synchronize with the beat of the metronome (see Methods, Eq. 3) and errors on this evolving target were penalized in the cost function, weighted by the parameter *δ* (Eq. 4). To match the average AFA exponent reported in the experimental data (Fig. 2), this parameter was set to 0.35 when gait was guided by a metronome. When walking freely, without any imposed cadence, this parameter was set to 0.

In the simulations presented below, the transition between these two values was linear and had a rise time of 1, 50, or 100 strides (Fig. 5A, green, yellow, and red, respectively). For each of these scenarios, we simulated 100 series of 700 strides and reported the mean evolution of the AFA exponent for all these series in Fig. 5B, using the same sliding-window analysis as for the experimental data. In the ON-OFF condition, only an instantaneous transition between *δ* = 0.35 and *δ* = 0 allowed us to reproduce the evolution of LRA observed experimentally, i.e., an immediate increase of the AFA exponent after the switching of the metronome. We performed the same segmented linear fit analysis as for the experimental data and found that the green trace, corresponding to an instantaneous switch of the control, started to increase seven strides after the first vertical line, compared to three strides for the experimental data. Thus, the almost instantaneous increase observed experimentally likely reflected an instantaneous change in the control strategy when the metronome was turned off.

**Figure 5:**
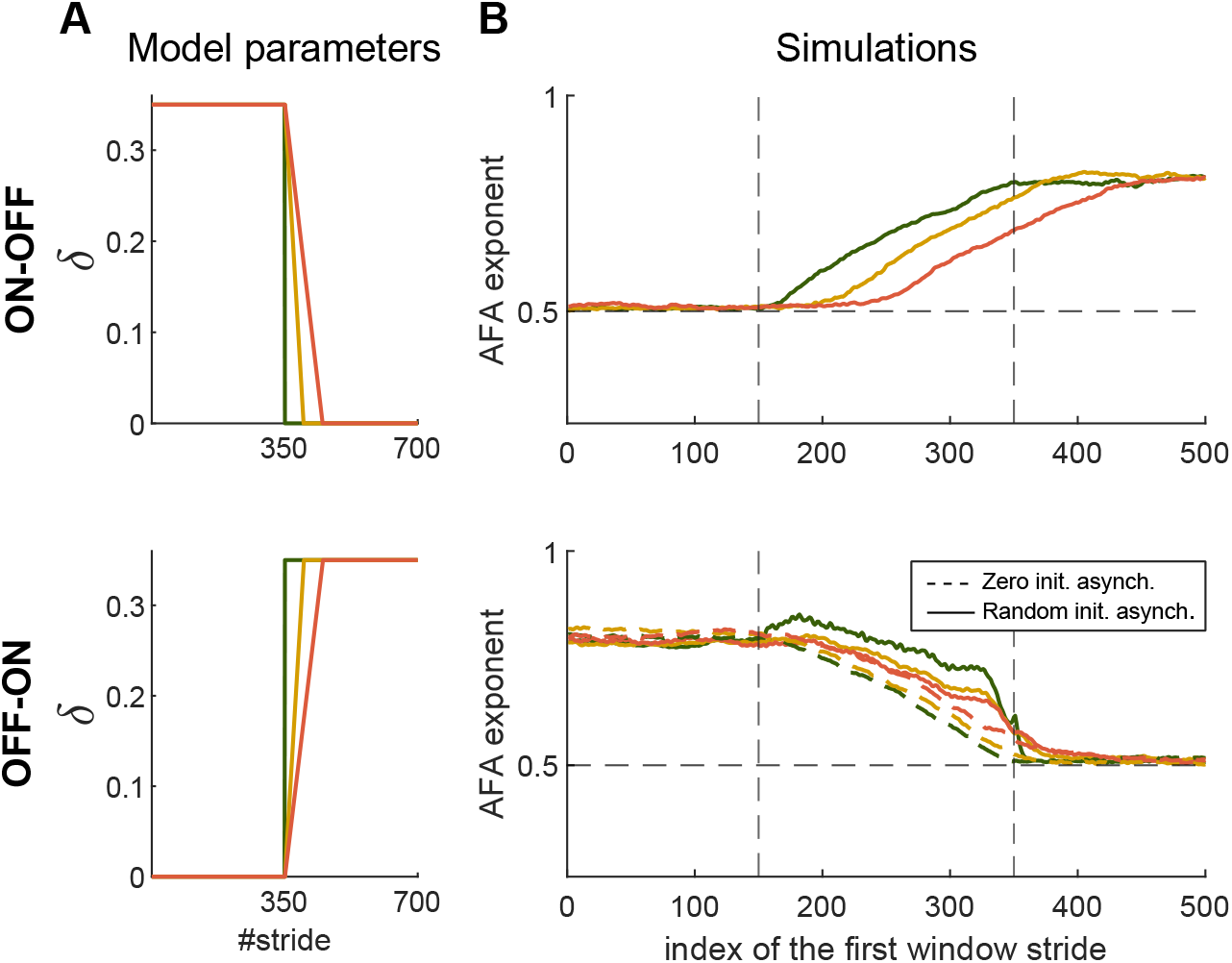
Transition analysis in simulations. (A) Value of parameter *δ* at each stride for three transion rates under study in ON-OFF (top) and OFF-ON (bottom) condions. (B) Evolution of the AFA exponent averaged over 100 simulated series. The colors indicate the parameters used to generate the stride series. In OFF-ON condition (bottom), the colored dotted traces correspond to an asynchrony with the metronome at the transition set to zero, solid lines correspond to a random initial asynchrony.

In the OFF-ON condition, we also manipulated the asynchrony at the transition when the metronome was turned ON. The three solid lines correspond to a random initial asynchrony, while the dotted lines correspond to an initial asynchrony of zero, i.e., perfect synchronization from the first stride. It appeared that a non-zero initial asynchrony, consistent with the experimental conditions, with an instantaneous transition in the parameter *δ* reproduced all features of the data. Among simulations including a random initial asynchrony, transitions at three different rates were tested. These three transitions exhibited an AFA exponent greater than 0.5 at the second vertical line, all three similar to experimental data (AFA exponent = 0.59 ± 0.046). Finally, only an instantaneous transition between *δ* = 0 and *δ* = 0.35 with initial random asynchrony reproduced the local increase in the AFA exponent occurring directly after the transition in the experimental data. To further understand the origin of this increase, we analyzed each simulation separately. We computed the difference between the maximum AFA exponent following the transition and the AFA exponent at the transition. We found a correlation coefficient of 0.57 between this variable and the initial asynchrony. The model thus suggested the following explanation for the transient increase reported in the OFF-ON transition: a random initial asynchrony, combined with immediate strong regulation, induced important deviations from the average pace for a few strides to synchronize with the metronome. This would yield locally changes in the time series and transient variations in the AFA, similarly to the effect of a sudden shift in mean value on the AFA exponent (Fig. 4).

We selected the parameters that best fitted the behavior observed experimentally, i.e., an instantaneous shift right after the switch in both conditions, and a random initial asynchrony in the OFF-ON condition. With these parameters, we simulated 20 new series for each condition. As we did with the participants’ data, we divided the series into two parts at the moment when the metronome was turned on/off (exactly in the middle of the series for the simulations) and compared their AFA exponents and standard deviations (Fig. 6).

**Figure 6:**
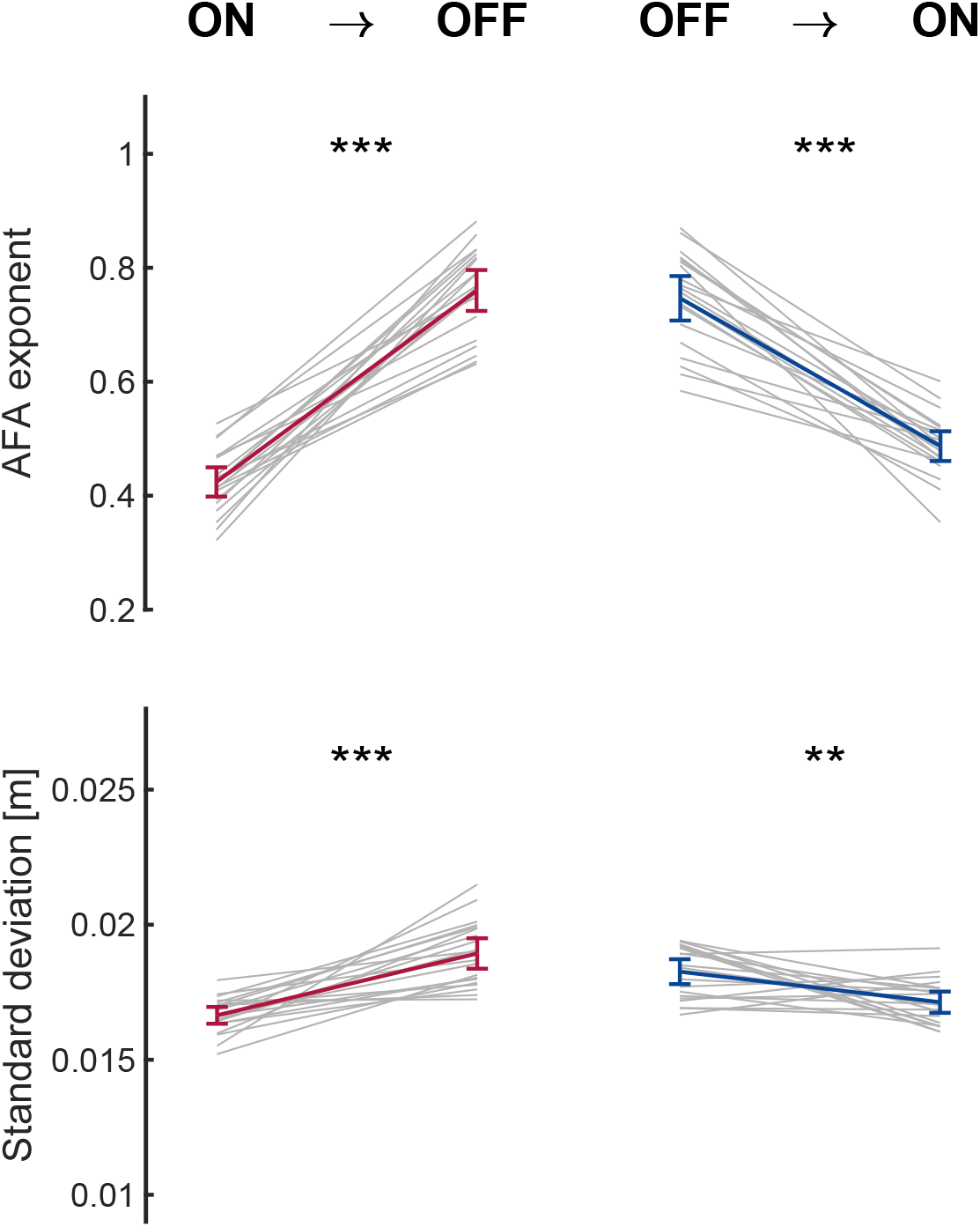
Analysis of differences between ON and OFF stride series in simulations. AFA exponent (A) and standard deviation (B) of simulated series of stride duraons before and after the switch of the metronome in ON-OFF condion (left) and OFF-ON condition (right). Group average and 95% confidence interval are shown in color and individual participants data are shown in gray lines. Stars indicate significant differences determined using paired, one-tail t-test: ** *p* < 0.01, *** *p* < 0.001.

The simulated series exhibited an average AFA exponent below 0.5 in ON parts of the series (0.42 ± 0.05 in ON-OFF condition), reproducing the anti-persistent stride series that appeared only when applying the AFA to longer series (here 350 strides), similarly to experimental data. The AFA exponent of the simulated series of stride duration was significantly impacted by the metronome (ON-OFF: Δ*AFA* = 0.34 ± 0.10, *t*(19) = 15.23, *p* < 10^−4^; OFF-ON: Δ*AFA* = −0.26 ± 0.09, *t*(19) = −13.14, *p* < 10^−4^), in a comparable way to the participants’ stride series (Fig. 6A). The effect sizes were *d* = 9.3 in the ON-OFF condition and *d* = −4.7 in the OFF-ON condition. We also observed a decrease in standard deviation when the metronome was activated in both conditions, as shown in Fig. 6B (ON-OFF: Δ*std* = −0.0018 ± 0.0014[*s*], *t*(19) = −7.11, *p* < 10^−4^; OFF-ON: Δ*std* = 1.15 ∗ 10^−3^ ± 0.0016[*s*], *t*(19) = 3.46, *p* = 0.0026). The effect sizes were *d* = 5.2 in the ON-OFF condition and *d* = −1.2 in the OFF-ON condition. Effect sizes on simulated data (both SD and AFA exponent) were larger than on experimental data, because inter-subject variability was not included in the model. However, the magnitudes of the ON and OFF differences were similar, and the effect was greater on the AFA exponent than on the SD, both in simulated and experimental data.

From the same 20 simulations, we extracted the series of asynchronies between heel strikes and metronome beats in the ON parts of the series. In accordance with the literature, these series displayed a strong persistence, with an average AFA exponent equal to 0.91 ± 0.059 (13, 18). The mean asynchrony was below 10^−4^[*s*] (≅ 0), and the standard deviation of the series was 0.0392 ± 0034 [*s*]. Since this variable was not measured in the protocol, we were unable to compare these values to experimental data. While the expected temporal structure of variability seemed successfully reproduced by the model, some authors demonstrated that people tend to adopt anticipatory strategies and strike before the tone (18, 19). They therefore displayed a negative mean asynchrony, something that we did not capture in our simulations.

We performed a sensitivity analysis to explore the impact of changes in the parameters. For *α* = *β* = 2.5, dividing or multiplying *δ* by 2 had a significant impact on the AFA exponent compared to *δ* = 0.35, but not on the standard deviation. When increasing this parameter, the regulation of the asynchrony was strengthened and the series of stride durations became more anti-persistent, with lower values of the AFA exponent. In contrast, a decrease of this parameter resulted in an increase of the AFA exponent with values above 0.5. Dividing *β* by 2 (*β* = 1.25) significantly increased the standard deviation of series of stride durations for *α* = 2.5 and *δ* = 0.35, but multiplying it (*β* = 5) had no impact both on the standard deviation and AFA exponent. By multiplying and dividing *α* by 2, the AFA exponent and standard deviation of stride durations did not change significantly, the effect was instead on the stride velocity, which is not of interest here. In addition, we assessed the impact of more minor changes in *δ*. Changing *δ* by ±10% had negligible effect on the standard deviation and the AFA exponent. Results were therefore robust to small changes in the cost function.

To summarize, the series of stride durations of participants exhibited anti-persistence and a slight decrease in standard deviation when they synchronized their gait with an isochronous metronome. When turning off the metronome in an ON-OFF condition, the AFA exponent rapidly increased to stabilize around a value similar to the self-paced condition, indicating the presence of LRA. In contrast, in the OFF-ON condition, the AFA exponent slightly increased shortly after the transition, then linearly decreased. These transitions were then reproduced in the model by an instantaneous change in the cost function for both conditions, provided that a random initial asynchrony was included in the model for the OFF-ON condition.

## Discussion

We investigated the time course of behavioral changes induced by synchronizing walking with a metronome. Indeed, we and other authors observed that synchronization of gait to an auditory stimulation profoundly modifies the statistical persistence of series of stride durations, with little effect on the mean and standard deviation of the same series. To characterize the transition from walking with to without a metronome and vice versa, 21 healthy young participants were asked to walk overground in three conditions, each lasting 15 minutes. The first condition was a control condition, with no constraints, to measure the participants’ natural pace. In the following two conditions, the metronome was activated either during the first or second half of the trial. We measured the evolution of the AFA exponents of these series over a sliding window and reproduced these traces in the framework of stochastic optimal control. In natural walking conditions (i.e., without a metronome), the model suggested that statistical persistence resulted from the selective and coordinated regulation of stride length and duration, while maintaining a constant target speed. Conversely, gait control when guided by a metronome was modelled by adding a cost regulating the asynchronies between the metronome beat and heel strike. This resulted in anti-persistent stride series and accounted for the changes observed in the standard deviation. This model allowed us to test several transitions between these two control modes and compare the resulting trends of AFA exponents with the experimental data. The results of the experiments were compatible with the simulations involving instantaneous switching of the control policy in both conditions, including an initial random asynchrony in the OFF-ON transition consistent with the experimental setting.

The model showed here that the emergence of anti-persistence in series of stride durations arises as a direct consequence of the presence of persistence in the series of asynchronies, which represent the integration of stride intervals. Although not directly measured in the experimental data, data from previous synchronization studies and simulated data consistently exhibited statistical persistence in asynchronies (13, 47). In the model, this was attributed to a flexible control mechanism (i.e., characterized by low control gains) acting on this variable, in a similar way to previous studies which attributed these statistical properties to a partial correction of asynchronies (22, 48). This is also analogous to the role of flexible control in the regulation of stride duration and length when walking freely on a treadmill in (23). In this study, they demonstrated that the level of control strongly determines the statistical persistence of the associated gait parameter. In particular, they showed that reducing control gains increased the level of persistence in series of stride durations, whereas over-correcting the errors in velocity resulted in anti-persistent series. In a recent study, higher control gains were associated with Parkinson’s disease, as this population often exhibits altered persistence, suggesting they were unable to exploit the task redundancy introduced when only a speed constraint is applied (43). Here we showed that anti-persistent series of stride durations arose not from over-correction of this gait parameter, but from partial correction of the integral of these series.

Furthermore, we suggest that humans rapidly adjusted the control policy, potentially within a single stride, to comply with the task requirements, which was followed by a gradual change in LRA. This conclusion is supported by the fact that the observed changes in the AFA exponent were consistent with simulations involving a rapid task-dependent control adjustment. This is all the more striking given that the two transitions tested here were of very different natures. They exhibited distinct patterns of change in the average AFA exponent, which could nevertheless be attributed to a rapid change in control within the model. In particular, we observed an unexpected increase in the AFA exponent at the activation of the metronome, then followed by a gradual decrease (Fig. 3C). This observation did not directly follow from a symmetrical switch in control strategy (simulations not shown). Yet, it was reproduced in the model by considering the fact that the initial asynchrony at the transition was random. This suggests that variations in LRA, both in amplitude and timing, could be fully explained by a very rapid task-dependent modulation of gait control.

Our interpretation emphasizes the role of rapid changes in control strategy without considering the effect of non-linear interactions that occur during gait control. While the non-linearities inherent to the biomechanics of walking are not necessary to explain the transitions, this does not rule out the possibility that they may contribute to the presence of persistence and anti-persistence. In that sense, Gates and colleagues showed that non-linearities were sufficient to explain LRA (21). We believe that both task-dependent control and complex biomechanics likely participate in the selection of efficient control strategies across individuals, and that the interaction between these factors remains to be investigated in more detail. It is possible to connect this model with continuous-time approaches (49, 50) and integrate it with musculoskeletal frameworks (51, 52). Such extensions would be valuable for characterizing the underlying neurophysiological mechanisms of cued walking in greater detail while taking biomechanical constraints into account.

The minimalistic model presented here, designed with a high level of abstraction, does not specify how different neural substrates implement or switch between control policies. Synchronizing movement with rhythmic auditory cues likely requires distributed processing across a large network, including sensorimotor cortex, supplementary motor area, premotor cortex, basal ganglia, and cerebellum (53, 54). The mobilization of these supraspinal structures may either supplement or override the automated regulation of free walking. Some authors argued that the disappearance of correlations under metronomic conditions could be interpreted as a complete overriding of the locomotor system’s dynamics by supraspinal influences (55). Conversely, others emphasized the fact that LRA were still present under metronomic conditions in the series of asynchronies, suggesting that the source of LRA was still at work but expressed differently (13). Regarding the transitions, our framework enforces abrupt transitions on a timescale corresponding to a stride (i.e., about 1s) to capture the observed behaviour. However, it does not specify how the nervous system detects a task change or triggers a switch in control policy. We hypothesize that the detection and triggering occur at a shorter timescale. For instance, the typical mean reaction time to the onset of an auditory stimulus is approximately 140 − 150*ms* (56). Here, we hypothesized that the introduction or removal of the metronome was perceived and used to change the control strategy almost instantaneously. Although evaluating the metronome frequency may require at least a couple of strides, our assumption is based on the fact that the auditory stimulus was salient and that the transition from ON to OFF required in the model a rapid switch. However, future extensions of the model could incorporate an additional module that estimates the transition less directly.

One of the main limitations of this study was the lack of assessment of the synchronization performance of participants through the measurement of the asynchronies between the heel strikes and metronome beats. The critical impact of these asynchronies on the evolution of LRA at the OFF-ON transition was unexpected, explaining why they were not included in the experimental design. Since this metric seems to play a central role in the control mechanisms, investigating its evolution in detail at the transitions could improve the interpretation of the evolution of AFA exponents. Synchronization performance might also directly impact the stride-to-stride variability (57). This could explain the slightly different results in the effect of the metronome on the standard deviation from one study to another.

It also remains unclear how the metronome interacts with the regulation toward the preferred pair of stride length and duration. This pair is partially defined by the biomechanical constraint of each subject and is associated with the gait parameters that minimize the energetic cost associated with the stride itself (58, 59). In free walking (without a metronome), there is a balance between the cost of the regulation (represented in the model by the term weighted by *γ* in Eq. 2) and the additional cost of a suboptimal pair (weighted here by *β*). Indeed, although this pair may be preferable in terms of energetic cost, it can also lead to significant changes from stride to stride, which are potentially costly. In this context, the metronome constraint often requires the adoption of different gait parameters from the preferred pair, thus introducing a conflict between these two terms of the cost function. The sensitivity analysis, used to test the impact of the parameter *β*, did not allow us to determine whether a reduction of this parameter occurred when the metronome was activated to mitigate this conflict. Indeed, this parameter has no impact on the statistical persistence of the stride duration under metronomic constraints (i.e., when *δ* = 0.35). However, it did have an impact on the variability of these same series. Participants may adopt different strategies in this regard, explaining the variation in synchronization performance and in the effect of the metronome on the stride-to-stride variability.

In this paper, we chose to study the impact of synchronization on rhythmic auditory cues with no variability. However, a growing body of literature advocates the use of fractal-structured cues, as they do not affect the level of persistence in gait variability among healthy adults (17). This type of cue could also restore healthy gait patterns in populations exhibiting a loss of complexity, such as older adults and patients with Parkinson’s disease (25–28). Although this could improve some biomarkers, there is no evidence that the mechanism underlying the recovery of statistical persistence with fractal cues is equivalent to the natural emergence of these patterns. In particular, in the model presented here, the persistence arose from the regulation of variability. This variability is assumed to originate from sensorimotor noise, which, in principle, is not correlated with an internal fractal timer. Therefore, although lower levels of persistence have been associated with an increased risk of falling (60, 61), the mechanism by which artificially restored persistence could improve gait control requires careful investigation. In addition, walking with invariant cues does not result in a loss of complexity, but rather in a shift of this complexity towards series of asynchronies. The model presented here offers an opportunity to investigate the use of fractal cues in the future. Like the type of cues (fractal-like, invariant, or random), two sensory modalities – auditory or visual – in a synchronization task can also affect gait parameters differently. In finger tapping synchronization tasks, it has been demonstrated that the persistence strength in the series of asynchronies depended on the sensory modality (62). These results seem to generalize to gait synchronization. In this regard, a study by Vaz and colleagues (63) showed that visual cues enhanced synchronization and had a distinct impact on step length persistence compared to auditory cues. Again, the model presented here provides an opportunity to investigate this further.

To conclude, this study highlights that transitions from self-paced walking to synchronization could result from instantaneous switches in control strategy, dictated by a cost function that exploits task redundancy when possible and regulates asynchronies when constrained. Our study offers a novel interpretation of the emergence of anti-persistence in series of stride durations when gait is guided by a metronome as the result of the regulation of strides to minimize asynchronies. Finally, this study proposes a new methodological framework that could be applied in future research to explore the use of cues in gait rehabilitation and its influence on gait control.

